# Nicotinamide N-Methyltransferase drives fibroblast activation and skin fibrosis in systemic sclerosis

**DOI:** 10.1101/2025.10.27.684733

**Authors:** Anela Tosevska, Karolina von Dalwigk, Peter Heil, Felix Kartnig, Ana Korosec, Birgit Niederreiter, Juan Manuel Sacnun, Thomas Köcher, Daniel Aletaha, Hans P. Kiener, Beate M. Lichtenberger, Leonhard X. Heinz, Thomas Karonitsch

**Author notes:** Corresponding author: Assoc. Prof. Dr. Thomas Karonitsch, Division of Rheumatology, Department of Medicine 3, Medical University of Vienna, Spitalgasse 23, 1090 Vienna, Austria.

## Abstract

**Background:** In systemic sclerosis (SSc), an autoimmune response leads to progressive fibrosis of the skin and internal organs, driven by aberrant activation of fibroblasts. The mechanisms dictating persistent dermal fibroblast (DF) activation and production of extracellular matrix (ECM) remain poorly understood. Nicotinamide N-methyltransferase (NNMT), a SAM-consuming enzyme that modulates cellular methylation potential, has been implicated in fibrotic tissue remodelling in metabolic and malignant diseases. Here, we identify NNMT as a key determinant in DF activation and fibrosis in SSc.

**Methods:** We analyzed bulk, single-cell RNA-Seq and spatial transcriptomics datasets from SSc skin. Functional studies were performed in TGFβ-activated primary human DFs using siRNA-mediated NNMT knockdown (KD) combined with RNA-Seq, metabolite profiling, ELISA, and western blotting. The role of NNMT-regulated transcription factors was assessed by QuantSeq 3′ RNA-Seq following ATF4, SOX9, or SRF KD.

**Findings:** NNMT was markedly upregulated in SSc skin and enriched in disease-expanded SFRP2⁺/COL8A1⁺ myofibroblast states. NNMT KD restored methylation balance by increasing the SAM/SAH ratio and H3K27me3 levels, and abrogated TGFβ-induced profibrotic programs regulating ECM production and collagen synthesis. Mechanistically, NNMT was required for TGFβ-induced upregulation of the transcription factors ATF4, SOX9, and SRF, which together orchestrate ECM gene expression and COL1A1 secretion.

**Interpretation:** These findings define a previously unrecognized TGFβ–NNMT-ATF4/SOX9/SRF axis that coordinates profibrotic transcriptional programs in DFs. Accordingly, NNMT functions as a central effector linking TGFβ signaling to DF activation and ECM remodelling. Targeting NNMT may thus represent a promising therapeutic strategy to attenuate skin fibrosis in SSc.

## Introduction

Systemic sclerosis (SSc) is a rare connective tissue disease characterized by vasculopathy, immune-mediated inflammation, and progressive fibrosis of the skin and internal organs (1, 2). The aetiology of SSc remains incompletely understood, although current models suggest that environmental triggers acting on genetically predisposed individuals initiate autoimmunity and vascular injury, which in turn promote fibroblast activation and progressive tissue fibrosis (3, 4). In early disease, immune- and endothelial-derived mediators such as transforming growth factor-β (TGFβ) trigger fibroblast activation, whereas epigenetic reprogramming results in sustained fibroblast-to-myofibroblast differentiation in later stages (5).

Despite recent advances in targeting inflammation, effective antifibrotic therapies for the skin are still lacking. Currently, no treatment halts or reverses dermal fibrosis, the most visible and debilitating feature of SSc. This therapeutic gap underscores the urgent need to identify novel molecular pathways that perpetuate dermal fibroblast (DF) activation and could be exploited for antifibrotic therapeutic intervention (6).

Nicotinamide N-methyltransferase (NNMT) catalyses the methylation of nicotinamide (NAM) using S-adenosylmethionine (SAM) as a methyl donor, producing 1-methylnicotinamide (1-MNA) and S-adenosylhomocysteine (SAH). Through consumption of SAM, NNMT modulates the intracellular SAM/SAH ratio, thereby regulating cellular methylation potential, epigenetic reprogramming, and gene expression (7–9).

NNMT has been implicated in a broad spectrum of metabolic disorders, including obesity, type 2 diabetes, and non-alcoholic fatty liver disease, where it contributes to energy imbalance, inflammation, and dysregulated lipid metabolism (10–13). NNMT is also highly expressed in several cancers and has emerged as a potential biomarker and metabolic driver of cancer progression (14). Recent studies link NNMT to cancer-associated fibroblast (CAF) activation and extracellular matrix (ECM) remodelling, highlighting its profibrotic potential (15). In line, experimental models of hepatic and renal fibrosis indicate that NNMT promotes fibrotic tissue remodelling (10, 16).

Based on its profibrotic functions in metabolic disease, liver cirrhosis, and CAF-mediated ECM remodelling, we hypothesized that NNMT acts as a key driver of DF activation and skin fibrosis in SSc. To probe into NNMT function in SSc, we analyzed publicly available transcriptomic datasets and found that NNMT is highly expressed in the skin of patients with SSc and in bleomycin-induced murine skin fibrosis. Further, we explored the significance of NNMT modulation on TGFβ-driven fibroblast activation and ECM production, in support of the hypothesis that NNMT inhibition may represent a novel antifibrotic strategy in SSc.

## Material and methods

### Analysis of publicly available datasets

Bulk RNA-Seq data from skin biopsies of SSc patients and HCs, including clinical metadata such as the mRSS, were obtained from published datasets (17). Raw FASTQ files were downloaded from SRA (SRP197303) and mapped to the human genome (GRCh38, GCF_000001405.39) using STAR v2.7.8a (18). Features were counted using HTSEq-count, and normalization and differential expression analysis were performed using DESeq2 v1.28.1 (19). In addition, we reanalyzed bulk RNA-Seq data from a bleomycin-induced murine model of dermal fibrosis (20). Raw counts were downloaded from GEO (GSE132869) and further processed with DESeq2. Signature enrichment was calculated using the “singscore” method from the hacksig package in R (21).

Single-cell RNA-Sequencing (scRNA-Seq) of skin was reanalyzed from publicly available datasets (22, 23). Data was downloaded from GEO (GSE138669 and GSE249279) and processed using Seurat v5.2.0 (24). For Tabib et al., cells with > 200 and < than 2500 features and < than 25% mitochondrial reads were retained (n=55465 cells). For Ma et al., cells with > than 200 features and 500 reads, as well as < than 15% mitochondrial content were retained (n=96157 cells). After log-normalization, clustering was performed using the top 20 PCs (GSE138669) or 30 PCs (GSE249279). Cell clusters were annotated based on marker genes from the FindAllMarkers in Seurat and verified against the corresponding publications. DFs were subsetted and further subclustered following the same standard Seurat workflow, resulting in 11478 and 24954 DFs, respectively. Cells from GSE138669 were clustered using 13 PCs. Due to a higher batch effect in GSE249279, samples were first integrated using CCA integration in Seurat, and clustering was performed based on 20 PCs. Gene signatures were calculated using AddModuleScore in Seurat and Plots were generated using scCustomize v3.0.1 package in R. Spatial transcriptomics data from Ma et al. were downloaded from GSE249279 and processed similarly using Seurat v5.3.0.

### Primary DF isolation and culture

Primary human DFs were isolated from abdominal skin obtained during plastic surgery with approval of the ethics committee (EC) of the Medical University of Vienna (EC Nr. 1695/2014). After removal of subcutaneous fat, skin was cut into strips and incubated with Dispase II (20 mg/mL; Roche) in PBS at 37 °C for 1 h to separate epidermis from dermis. The dermis was minced and digested for 1 h at 37°C in DMEM/10% FCS containing Collagenase I (50 mg/mL; Gibco), Collagenase II (50 mg/mL; Gibco), Collagenase IV (50 mg/mL; Sigma), Hyaluronidase (10 mg/mL; Sigma), and DNase I (5 mg/mL; Sigma). The cell suspension was filtered, washed, and plated in DMEM + 10% FCS (GIBCO) and 1% penicillin–streptomycin (GIBCO). DFs were maintained with weekly medium changes and passaged upon confluence. DFs between passages 5–10 were used for experiments.

For TGFβ stimulation DFs were starved in DMEM with 2% FCS for five hours and treated with 20 ng/ml recombinant human TGFβ (R&D) for up to 72 hrs.

### siRNA-mediated knockdown

DFs were transfected with pooled siRNAs (ON-TARGETplus SMARTpool, Dharmacon) targeting NNMT (L-010351-00-0005), ATF4 (L-005125-00-0005), SOX9 (L-021507-00-0005), or SRF (L-009800-00-0005), or with non-targeting control siRNA, using Lipofectamine RNAiMAX (Thermo Fisher) as previously described (25). KD efficiency was confirmed by western blotting.

### ELISA

ELISA for COL1A1 was performed using a commercial ELISA kit (R&D) according to the manufacturerś instructions.

### Western blot

DFs were lysed in RIPA buffer (Thermo Fisher) supplemented with protease/phosphatase inhibitors (SIGMA). Equal amounts of protein were separated by SDS-PAGE and transferred to nitrocellulose membranes. Membranes were probed with ABs against SPARC, POSTN, ACTA2/αSMA, LOXL2, SOX9, SRF, ATF4, H3, H3K4me3, H3K27me3 (Cell Signaling Technology), and NNMT (Atlas Antibodies and Cell Signaling). Tubulin (Sigma) or Actin (Sigma) served as loading controls. HRP-conjugated secondary ABs (Cell Signaling Technology) and Clarity ECL substrates (Bio-Rad) were used for detection on a ChemiDoc Imaging System (Bio-Rad). When required, membranes were stripped using ReBlot Plus Strong solution (EMD Millipore).

### High-content microscopy and computational image analysis

DFs were seeded onto 96-well glass-bottom plates for 24 hours, fixed with 4% paraformaldehyde and permeabilized with 0,3% Triton X-100. The actin cytoskeleton was stained with Alexa Fluor 488– conjugated phalloidin (Thermo Fisher), and nuclei were counterstained with DAPI (Sigma). Images were acquired using an Opera Phenix high-content screening system (PerkinElmer). Quantitative analysis of cell morphology (cell area, compactness, eccentricity and perimeter) and actin intensity and distribution was performed using CellProfiler (v4.2.8) and data were plotted in R (v4.5.0)

### Metabolite analysis

Metabolites were extracted from DFs using methanol:acetonitrile:water (2:2:1, v/v/v) following three freeze–thaw cycles in liquid nitrogen. Extracts were incubated at −20 °C for one hour and centrifuged at 16,000 × g for 15 min at 4 °C. Supernatants were stored at −80 °C until analysis. Metabolites were analyzed with hydrophilic interaction liquid chromatography (HILIC) LC–MS/MS (liquid chromatography tandem mass spectrometry) and reversed phase LC-MS/MS. RP-LC-MS/MS was performed by employing an Ultimate 3000 HPLC system, coupled to a TSQ Altis mass spectrometer (both Thermo Fisher Scientific). Briefly, dried biological samples were resolubilized by adding 1% formic acid in water and 1 µl of each sample was injected onto a Kinetex (Phenomenex) C18 column (100 Å, 150 x 2.1 mm) connected with a guard column, and employing a 4-minute-long linear gradient from 97% A (1 % acetonitrile, 0.1 % formic acid in water) to 60% B (0.1 % formic acid in acetonitrile) at a flow rate of 80 µl/min. Employing selected reaction monitoring (SRM) in the positive ion mode, 1-methylnicotinamide was quantified with the SRM-transition m/z 137.1 to m/z 94. In HILIC-MS/MS, an Ultimate 3000 HPLC system directly coupled to a TSQ Quantiva mass spectrometer (both Thermo Fisher Scientific) was used. 1 µl of each sample was directly injected onto a polymeric iHILIC-(P) Classic HPLC column (HILICON, 100 x 2.1 mm; 5 µm) connected with a guard column. A flow rate of 100 µl/min was used and a linear gradient (A: 85% acetonitrile, 10% 10 mM aqueous ammonium bicarbonate, 5% 10 mM aqueous ammonium bicarbonate; B: 10 mM aqueous ammonium bicarbonate) was applied. The linear gradient started with 6% B, increasing to 80% B in 17 minutes. The following SRM transitions were used for quantitation in the positive ion mode: m/z 399.1 to m/z 250.1 (S-adenosylmethionine), m/z 385 to m/z 136 (S-adenosylhomocysteine). In both assays, retention times, SRM-transitions, and optimal collisional energies were determined by authentic standards. All data interpretation was performed using Xcalibur (Thermo Fisher Scientific).

### Real-time quantitative polymerase chain reaction

Total RNA was isolated using the RNeasy purification kit (Qiagen). RNA concentration and integrity was assessed using a Nanodrop spectrophotometer. cDNA was synthesized using the Omniscript kit (Qiagen) and amplified with SYBR Green I Master Mix (Roche) on a LightCycler system (Roche). Relative expression was calculated by the 2−ΔΔCt method, using GAPDH as reference. Primer sequences: NNMT—forward TGGCTTCTGGAGGAAAGAGA, reverse AATCAGCAGGTCTCCCTTCA; GAPDH—forward TGATGACATCAAGAAGGTGGTGAAG, reverse TCCTTGGAGGCCATGTGGGCCAT.

### RNA-Seq library preparation

Total RNA was isolated using the RNeasy purification kit (Qiagen) and quantified using the Qubit® RNA BR Assay Kit (Thermo Fisher Scientific). The RNA integrity number (RIN) was determined using the RNA 6000 Pico Kit (Agilent) on a 2100 Bioanalyzer High-Resolution Automated Electrophoresis instrument (G2939A/B, Agilent). RNA-Seq libraries were prepared with the NEBNext® Poly(A) mRNA Magnetic Isolation Module E7490, the NEBNext® Ultra™ II Directional RNA sample preparation kit E7760 and the NEBNext® Multiplex Oligos for Illumina (Unique Dual Index UMI Adaptors RNA) (New England Biolabs). NGS library concentrations were quantified with the Qubit® 1X dsDNA HS Assay Kit (Thermo Fisher Scientific), and the size distribution was assessed using the High Sensitivity DNA Kit (Agilent) on a 2100 Bioanalyzer instrument. Before sequencing, sample-specific NGS libraries were diluted and pooled in equimolar amounts.

### QuantSeq3’RNA-Seq library preparation

NGS libraries were prepared from total RNA samples with the QuantSeq 3’ mRNA-Seq V2 Library Prep Kit with the UDI kit (Lexogen). Resulting library concentrations were quantified with the Qubit® 1X dsDNA HS Assay Kit (Thermo Fisher Scientific) and the fragment size distribution was assessed using the High Sensitivity DNA Kit (Agilent, Santa Clara) on a 2100 Bioanalyzer High-Resolution Automated Electrophoresis instrument (Agilent). Before sequencing, sample-specific NGS libraries were diluted and pooled in equimolar amounts.

### Next-generation sequencing and raw data acquisition

Expression profiling libraries were sequenced on a HiSeq 3000 or NovaSeq 6000 instrument (Illumina) following 50-base-pair, single-end, 100-base-pair paired-end or 100-base-pair single-end recipe.

### Transcriptome analysis

NGS reads were mapped to the Genome Reference Consortium GRCh38 assembly via “Spliced Transcripts Alignment to a Reference” (STAR, 2.7.9a) utilising the “basic” GENCODE/Ensembl transcript annotation from versions e100 (April 2020), v43 (February 2023) and v46 (May 2024) as reference transcriptome. Metadata annotation, such as the start of NGS adapter sequences (in XT tags), or unique molecular index (UMI) sequences (in RX tags) and UMI base quality scores (in QX) tags were propagated from unaligned BAM files to the aligned BAM files via Picard MergeBamAlignment and duplicated alignments were marked with Picard MarkDuplicates in a UMI-aware manner but not removed. Features were counted with the Bioconductor GenomicAlignments (1.34.0) package via the summarizeOverlaps function in Union mode, ignoring secondary alignments, alignments not passing vendor quality filtering and for UMI-supporting protocols only, also duplicate alignments (26). Gene counts were normalized using DESeq2 v1.48.1. Differential expression was defined as an absolute log2 fold change >1 with a Benjamini–Hochberg adjusted P-value <0.05 after filtering for mean normalized counts >10 (RNA-Seq) or counts >5 (Quantseq 3’ RNA-Seq). Volcano plots were generated using EnhancedVolcano (v1.26.0, R package). Plots were constructed in R v4.5.1 using ggplotv4.0.0, ggpubr v0.6.1 and pheatmap v1.0.13 and ComplexUpset v1.3.3.

### Enrichment analysis and enrichment term clustering

Differentially expressed genes and gene signatures were functionally enriched, clustered and summarized using the SummArIzeR tool in R. Databases used: “BioPlanet_2019”, “GO_Biological_Process_2023”, “Reactome_2022”, and “MSigDB_Hallmark_2020” from enrichR (27). The top 10 terms based on adjusted p-value per database and comparison were selected and terms with < than 5 genes were filtered out. Similarity scores were calculated based on Jaccard similarity, and clusters were inferred from the similarity matrix using the Walktrap community finding algorithm. Edges < 0.18 (figure 2F and 4E) and < 0.2 (figure 3E), were excluded.

### Gene co-expression analysis and network construction

The siRNA datasets (siNNMT, siATF4, siSOX9 and siSRF) were merged, normalized using DESeq2 and batch-corrected using limma v3.64.3 (28). Co-expression analysis was performed using CEMiTool v1.32.0 with the following parameters: network_type = “signed”, tom_type = “signed”, merge_similar = F, min_ngen = 20, diss_thresh = 0.9. The resulting modules were inspected for gene membership and the module containing NNMT (M10) was used to intersect with the union of all genes from fig 4D (TGF-upregulated siGenes-downregulated signatures). The resulting 33 genes were used as an input for GeneMANIA (29) using all coexpression datasets as input with automatically selected weighing method. The resulting network was exported and visualized with Cytoscape v3.10.3.

### Statistical analysis

Data are presented as mean ± SED unless otherwise indicated. Statistical analyses were performed using GraphPad Prism (v10) or R. For normally distributed data, paired or unpaired t-tests were applied; for non-normally distributed data, Mann–Whitney U or Wilcoxon signed-rank tests were used. Statistical significance was defined as P < 0.05.

### Data Availability

The RNA-Seq/Quant 3’ Seq data generated in this study will be available in the ArrayExpress database (https://www.ebi.ac.uk/arrayexpress) upon publication.

## Results

### NNMT is upregulated in SSc skin and enriched in disease-expanded fibroblast states

Analysis of publicly available bulk RNA-Seq data (17) showed significantly higher NNMT expression in the skin from patients with SSc compared to healthy controls (HCs) (Figure 1A). NNMT transcript levels positively correlated with the modified Rodnan skin score (mRSS), a validated clinical measure of skin fibrosis (30) (Figure 1B and 1C). To substantiate these observations, we explored NNMT expression in a publicly available dataset of bleomycin-induced dermal fibrosis in mice (20), a widely used preclinical model of SSc (31) and found a significant upregulation of NNMT in fibrotic skin compared to controls (Figure 1D), consistent with the results observed in patients.

**Figure 1.**
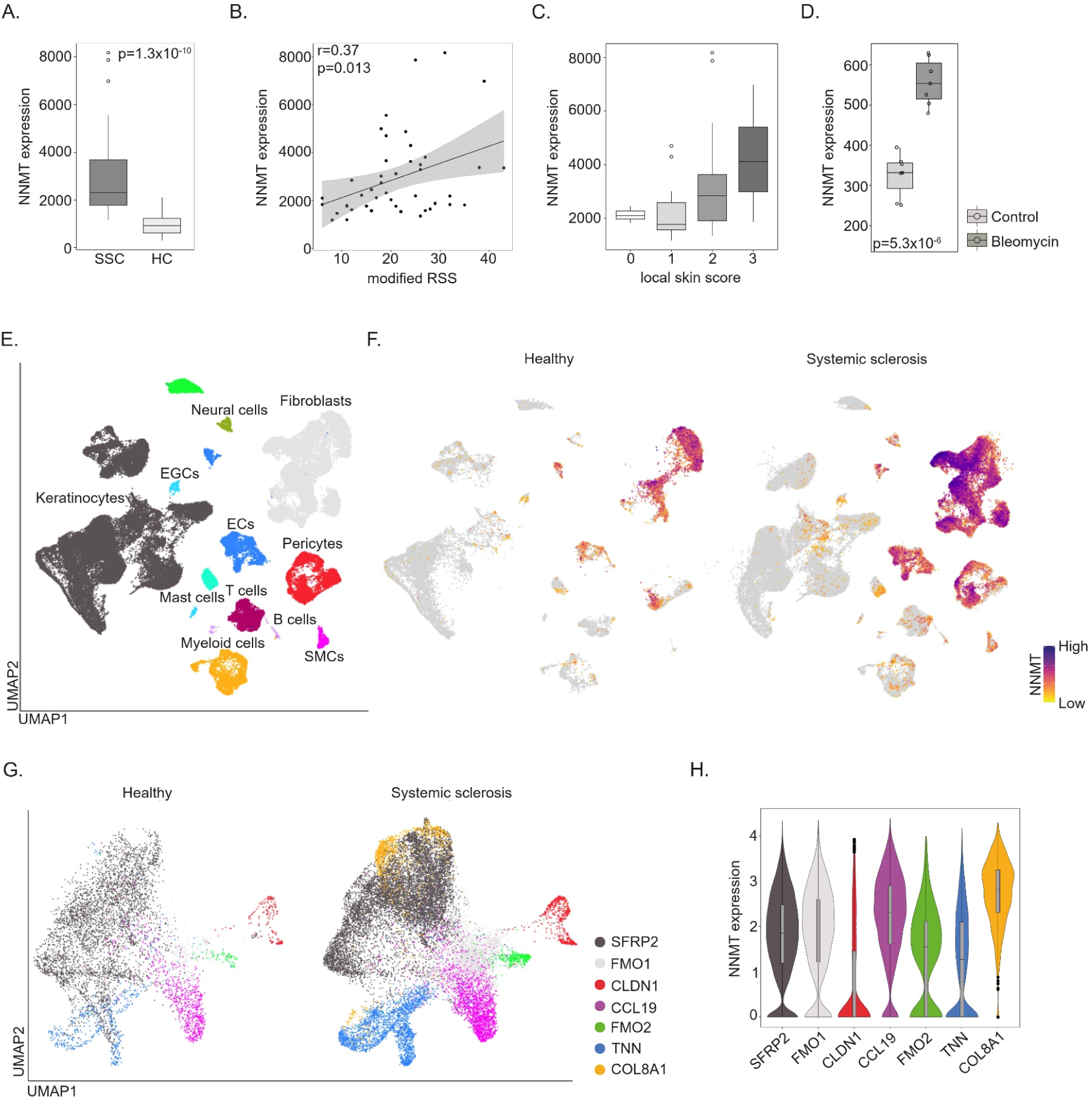
NNMT is upregulated in SSc skin and enriched in disease-expanded fibroblast states. **A.-C.** Analysis of publicly available bulk RNA-Seq data from the Prospective Registry for Early Systemic Sclerosis (PRESS) cohort (17). **A.** NNMT mRNA expression in skin biopsies from patients with SSc (n=47) and healthy controls (HCs; n=32). **B.** Correlation between NNMT expression and the modified Rodnan Skin Score (mRSS, n=44). **C.** Association between NNMT and the local skin score at the biopsy site (n=44). **D.** NNMT mRNA expression in skin samples from mice treated with bleomycin or vehicle control, as assessed by bulk RNA-Seq (20) (n=8). **E-H.** Analysis of publicly available scRNA-Seq data from skin biopsies of SSc (n=22) patients and HCs (n=18) (23). **E.** UMAP plot showing major skin cell populations. **F.** UMAP plots showing NNMT expression (log2 normalized counts) across cell types in HC and SSc. **G.** UMAP plot showing DF subtypes identified in HC and SSc. **H.** Violin plots showing NNMT expression levels (log2 normalized counts) across DF subtypes. For all experiments, n refers to the number of independent donors (biological replicates).

Given the cellular heterogeneity of DFs revealed by scRNA-Seq (32, 33), we reanalyzed two skin scRNA-Seq datasets from HCs and SSc patients (22, 23). NNMT expression was minimal or absent in keratinocytes and immune cells, and mostly confined to stromal compartments, including endothelial cells, pericytes, and most prominently DFs (Figure 1E and 1F; Supplementary Figure 1A and 1B). Reclustering of DFs recovered subtype structures consistent with the published data (Figure 1G; Supplementary Figure 1C). Although NNMT was expressed across multiple DF states, the highest levels were observed in SSc-enriched populations, specifically COL8A1+ DFs (Figure 1H) and SFRP2/PRSS23+ DFs (Supplementary Figure 1D), which were absent in HC skin. These two populations are transcriptionally related, representing overlapping states within the broader SFRP2+ DF lineage, and correspond to myofibroblast precursors or differentiated myofibroblasts (22, 23). In addition, NNMT was also highly expressed in the CCL19+ DF subtype, a population previously described to interact closely with immune cells (33). Thus, NNMT is not uniformly expressed across fibroblast states, but is preferentially enriched in distinct, disease-associated CCL19+ and SFRP2+ subsets that expand in SSc and align with clinical measures of skin disease severity (34), positioning NNMT within the SSc myofibroblast lineage.

### NNMT loss rewires DF methylation, cytoskeleton, and ECM programs

To define the (patho)physiological role of NNMT, we isolated human DFs and performed siRNA-mediated knockdown (KD) using pooled siRNAs targeting NNMT (25) (Figure 2A). As reported in CAFs (15, 35), 1-MNA, the enzymatic product of NNMT, was depleted in NNMT-KD DFs (Figure 2B). Metabolite profiling further revealed an increase in the SAM/SAH ratio in NNMT-KD cells (Figure 2B), indicating an elevated methylation potential. Consistently, global H3K27me3 levels were increased upon NNMT KD (Figure 2C), confirming on-target metabolic and chromatin-related effects. NNMT KD also induced marked morphological changes: NNMT-KD DFs were smaller, lost the typical spindle-shaped morphology, and displayed pronounced actin cytoskeleton reorganization (Figure 2D; Supplementary Figure 2).

**Figure 2.**
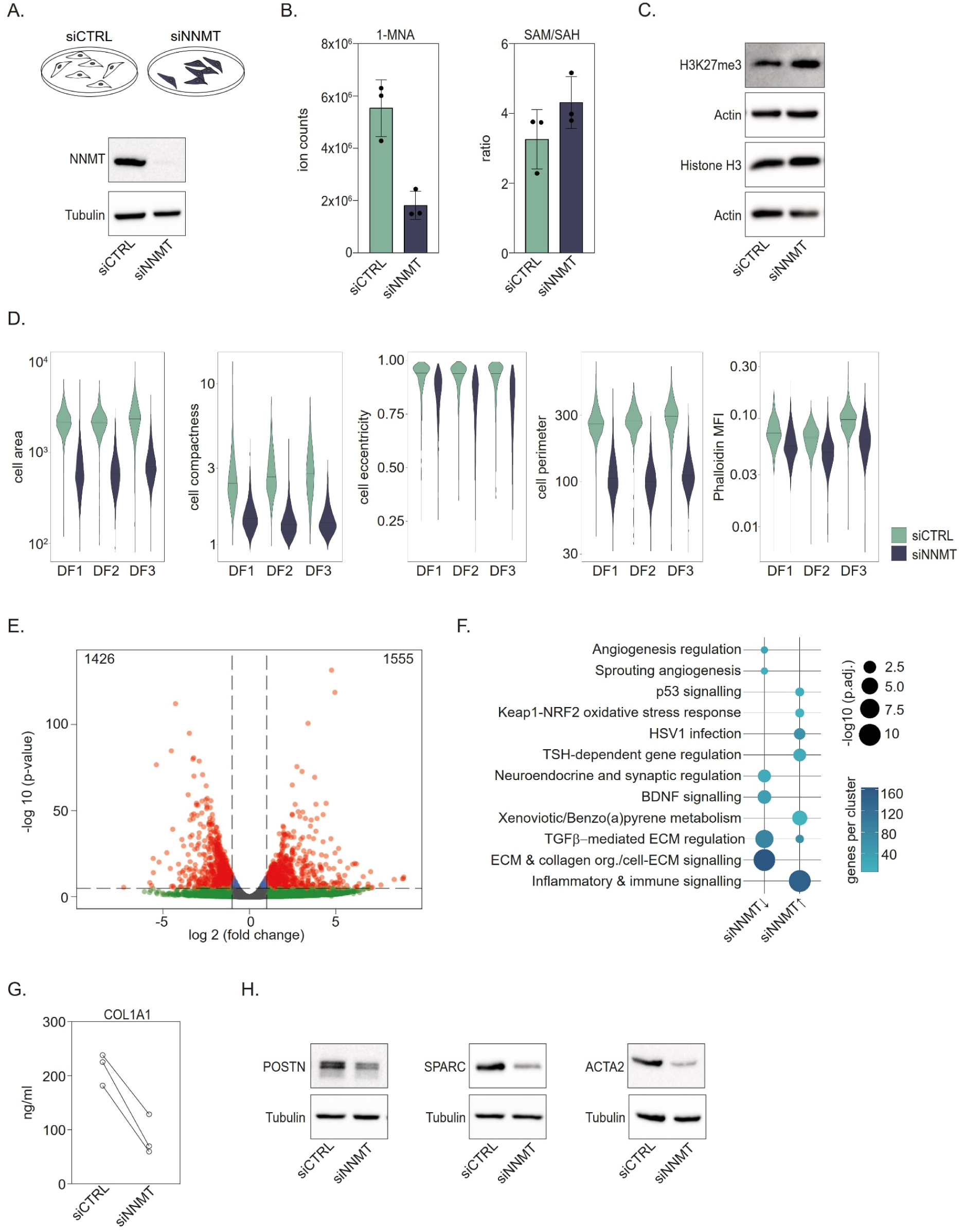
NNMT loss rewires DF methylation, cytoskeleton, and ECM programs. **A.-H.** Human DFs were transfected with control or NNMT-targeting siRNA pools. **A.** Schematic of the experimental setup and KD efficiency as assessed by western blots. **B.** Metabolite analysis was performed by reverse-phase liquid chromatography-tandem mass spectrometry (RP-LC-MS/MS). Shown are levels (ion counts) of 1-methylnicotinamide (1-MNA) and the SAM/SAH ratio (ion counts). Mean±standard deviation (n=3). **C.** Representative western blots showing expression of H3 and H3K27me3 (n=3). **D.** Cell morphology and actin intensity of DFs from three different donors (DF1, DF2 and DF3). Violin plots show cell area, compactness, eccentricity and perimeter as well as phalloidin mean fluorescence intensity (MFI) in siControl (siCTRL) and siNNMT conditions. **E.-F.** RNA sequencing of siCTRL and siNNMT DFs (n=3). **E.** Volcano plot illustrating the effect of NNMT KD on the DF transcriptome (log_2_ fold change> 1; adjusted p < 0.05). **F.** Functional enrichment analysis of genes differentially regulated in NNMT KD DFs. **G.** COL1A1 secretion measured by ELISA (n=3). **H.** Representative western blots showing expression of POSTN, SPARC and ACTA2 (n=3). For all experiments using DFs, n refers to the number of independent donors (biological replicates).

RNA-Seq of NNMT-KD DFs uncovered widespread transcriptomic reprograming (Figure 2E), with 1,556 genes upregulated and 1,426 downregulated relative to control siRNA. Functional enrichment analysis identified coordinated downregulation of processes related to ECM remodeling and collagen organization (Figure 2F). Accordingly, transcripts encoding ECM structural proteins (COL1A1, COL3A1, FN1), ECM organizers (POSTN, SPARC, THBS1/2/4), and DF activation markers (ACTA2/αSMA) were reduced in NNMT-KD DFs. To independently validate these findings using a complementary approach, we reanalyzed publicly available RNA-Seq data from mouse 3T3 cells overexpressing NNMT (15). Strikingly, genes downregulated by NNMT KD in our dataset were conversely upregulated upon NNMT overexpression (Supplementary Figure 3), supporting a reciprocal transcriptional relationship. These transcriptional changes were validated at the protein level by ELISA for COL1A1 (Figure 2G) and immunoblotting for SPARC, POSTN and ACTA2 /αSMA (Figure 2H).

### NNMT is TGFβ-inducible and required for full execution of profibrotic programs

TGFβ is a central driver of fibroblast activation and myofibroblast differentiation in SSc (4). We therefore examined whether NNMT is TGFβ-responsive and whether it is necessary for the full execution of TGFβ-driven profibrotic programs. qPCR (Figure 3A) and western blot (Figure 3B) showed that TGFβ stimulation upregulated NNMT in DFs. In line, analysis of publicly available RNA-Seq data from bleomycin-induced skin fibrosis (20) showed that pharmacologic blockade of TGFβ receptor I with SB525334 prevented NNMT induction (Supplementary Figure 4), indicating that NNMT lies downstream of TGFβ signaling.

**Figure 3.**
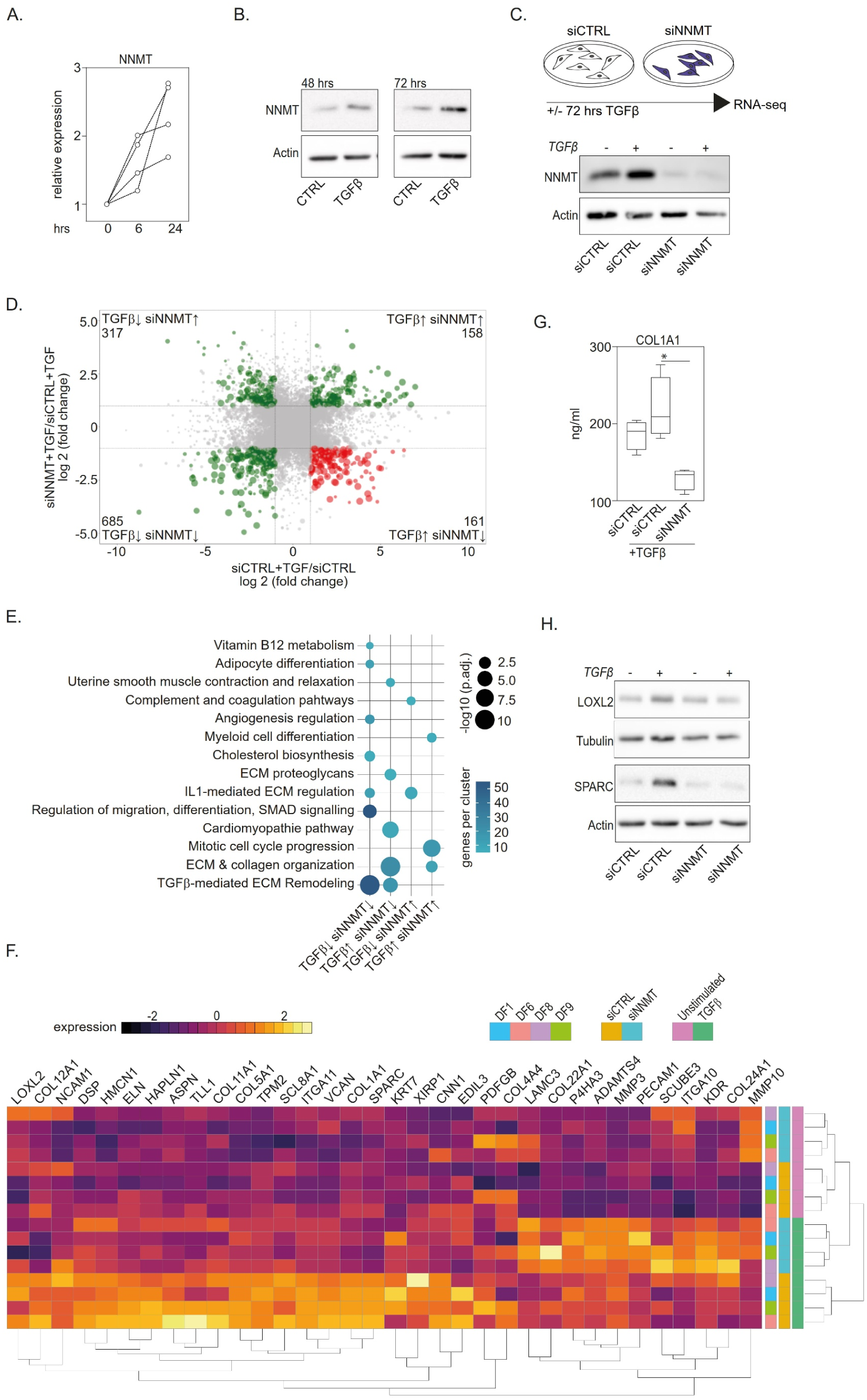
NNMT is TGFβ-inducible and required for full execution of profibrotic programs. **A.** qPCR of NNMT expression in DFs from four independent donors stimulated with TGFβ (20 ng/ml). Expression is shown relative to unstimulated cells and GAPDH. **B.** Representative western blots of DFs (n=4) stimulated with TGFβ (20 ng/ml) for the indicated time points. **C.** Schematic of the experimental setup of RNA-Seq workflow, and KD efficiency in DFs stimulated with TGFβ (20 ng/ml) for 72 hours, as assessed by western blots. **D.** Scatter plot illustrating the influence of NNMT KD on TGFβ-regulated genes, as measured by RNA-Seq (n=4). Genes highlighted in red (lower right quadrant) represent TGFβ-induced genes whose induction is blunted by NNMT KD. These genes include COL1A1, LOXL2 and SPARC (cutoffs: log_2_ fold change>1; adjusted p<0.05). **E.** Gene set enrichment analysis of TGFβ-regulated genes that are affected by NNMT KD. **F.** Heatmap showing genes enriched in the ECM and collagen organization pathways. **G.** COL1A1 secretion measured by ELISA. DFs from four different donors were stimulated for 72 hrs with TGFβ (20 ng/ml) (n=4, Paired t-test). **H.** DFs were stimulated with TGFβ (20 ng/ml) for 72 hours. Representative western blots (n=3) showing the expression of LOXL2 and SPARC. For all experiments using DFs, n refers to the number of independent donors (biological replicates).

To determine NNMT’s functional contribution to downstream TGFβ gene programs, we performed RNA-Seq on NNMT-KD DFs following TGFβ stimulation (Figure 3C-F). NNMT loss significantly altered the expression of multiple TGFβ-target genes (Figure 3D). Functional enrichment analysis revealed a prominent attenuation of well-known TGFβ-driven profibrotic programs, including ECM and collagen organization as well as remodeling (Figure 3E). Exemplary TGFβ-inducible genes closely linked to SSc, including COL1A1, COL5A1, LOXL2, SPARC (Figure 3F), and the myofibroblast marker ACTA2/αSMA (not shown) were significantly reduced in NNMT-KD DFs relative to control siRNA. Concordant protein-level validation showed decreased COL1A1 (Figure 3G) and reduced LOXL2 and SPARC (Figure 3H) in NNMT-KD DFs upon TGFβ stimulation.

Together, these data position NNMT as a TGFβ-inducible effector that is required for the full transcriptional execution of profibrotic programs in DFs.

### NNMT couples TGFβ signaling to a transcription-factor triad that drives profibrotic programs

To pinpoint transcriptional effectors downstream of NNMT, we identified NNMT-dependent, TGFβ-modulated transcription factors (TFs). A heatmap of significantly affected TFs highlighted ATF4, SOX9, and SRF as promising candidates (Figure 4A). At the protein level, TGFβ increased ATF4, SOX9 and SRF, and this induction was prevented by NNMT KD (Figure 4B), placing these factors downstream of the TGFβ-NNMT axis. We next profiled TGFβ-stimulated DFs after ATF4, SOX9, or SRF KD using QuantSeq3’ RNA-Seq (Figure 4C). Per-condition differential expression analysis showed that several TGFβ-modulated genes were downregulated following each KD (Supplementary Figure 5). An UpSet analysis integrating these data with the TGFβ-dependent NNMT KD gene signature, revealed a notable intersection of TGFβ-upregulated genes attenuated by all KDs (Figure 4D), indicating convergent control of a shared TGFβ-regulated transcriptional module. Consistently, functional enrichment analysis of the TGFβ-upregulated gene set across KD conditions uncovered attenuation of canonical profibrotic programs, including ECM and collagen organization as well as integrin and growth factor signaling in cell adhesion and TGFβ-mediated ECM regulation (Figure 4E). This transcriptional attenuation translated into reduced ECM output, as TGFβ-induced COL1A1 secretion was significantly diminished following ATF4, SOX9, or SRF KD (Figure 4F).

**Figure 4.**
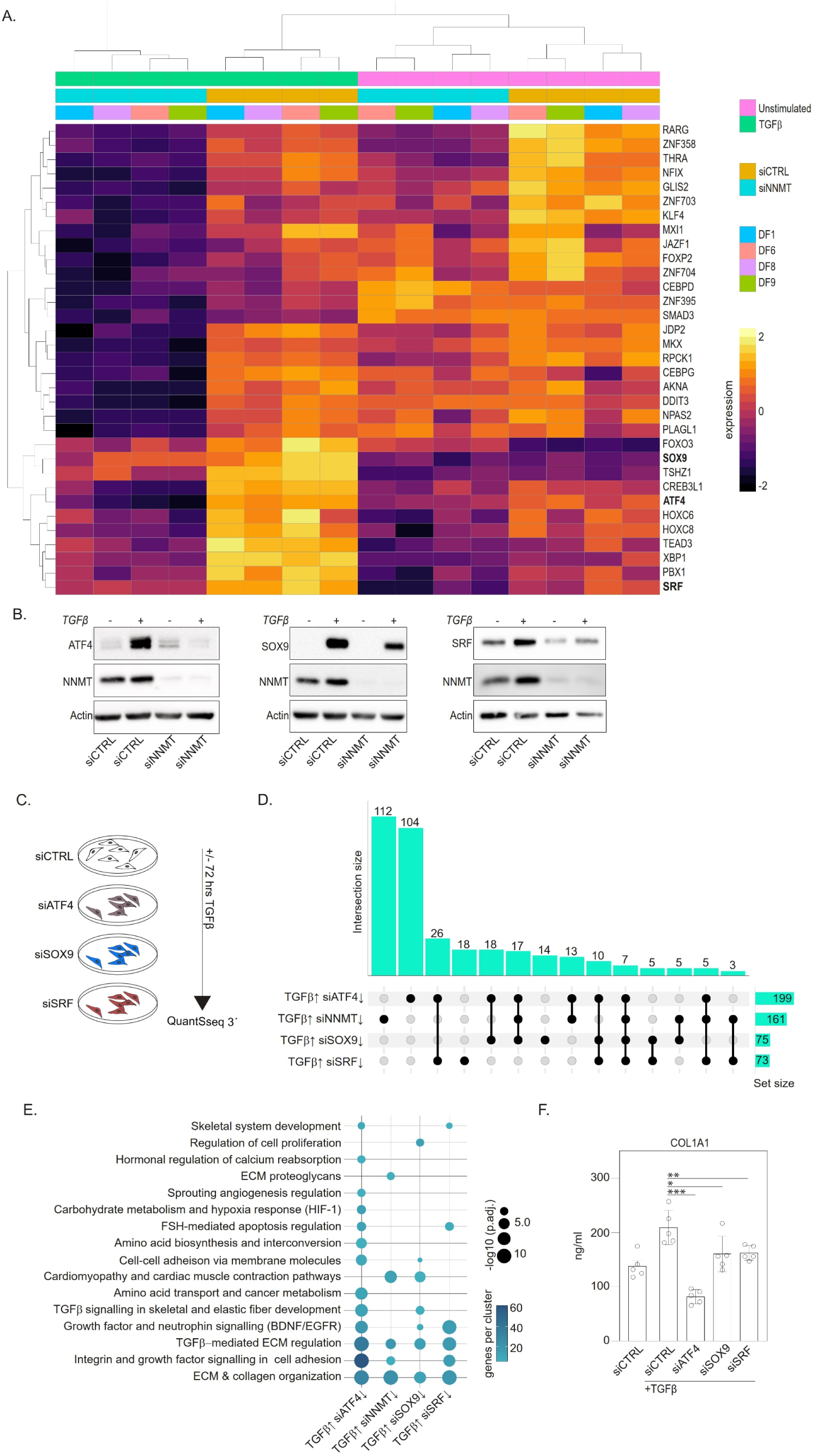
NNMT couples TGFβ signaling to a transcription-factor triad that drives profibrotic programs. **A.** Heatmap showing NNMT-regulated transcription factors that are modulated by TGFβ, as assessed by RNA-Seq. **B.** Representative western blots (n=3) showing the expression of ATF4, SOX9 and SRF in DFs transfected with siCTRL or siNNMT and stimulated with TGFβ (20 ng/ml; 72 hours). **C.-E.** DFs from five independent donors were transfected with non-targeting siRNA or siRNA pools targeting ATF4, SOX9 or SRF. mRNA was analyzed by QuantSeq 3’ RNA-Seq. **C.** Schematic representation of the experimental setup. **D.** Upset plot showing the overlap between TGFβ-upregulated genes whose induction is impaired by knockdown of ATF4, NNMT, SOX9 or SRF. **E.** Functional enrichment analysis of TGFβ-upregulated genes that are impaired by the knockdown of ATF4, NNMT, SOX9, or SRF. **F.** COL1A1 ELISA of supernatants from DFs that were transfected with control siRNA or siRNA pools targeting ATF4, SOX9 or SRF and then stimulated with TGFβ (20 ng/ml; 72 hours) (n=5, Paired t-test). Mean±standard deviation. For all experiments using DFs, n refers to the number of independent donors (biological replicates).

Together, these findings support an NNMT-dependent ATF4/SOX9/SRF regulatory axis that is required for broad execution of TFGβ-driven profibrotic programs in DFs.

### The TGFβ-NNMT-ATF4/SOX9/SRF network localizes to disease-expanded myofibroblasts lineage and niches in SSc

Using the scRNA-Seq dataset introduced in Figure 1, we mapped NNMT and its downstream TFs across DF states. NNMT, ATF4, SOX9, and SRF displayed similar spatial and transcriptional expression patterns, with the strongest overlapping signals in SFRP2+ DFs, and particularly in COL8A1+ myofibroblasts (Figure 5B). Consistent with their profibrotic role, both the collagen gene and ECM module scores were highest in DF states expressing high levels of NNMT, ATF4, SOX9, and SRF. To test whether the in-vitro dependencies identified for NNMT (Figure 3) and its downstream transcription factors (Figure 4) converge within specific DF states in vivo, we projected TGFβ-dependent KD gene signatures onto the same scRNA-Seq data to visualize their enrichment across DF subpopulations. TGFβ-upregulated genes were enriched within a subset of COL8A1+ DFs, and TGFβ-induced genes attenuated by NNMT, ATF4, SOX9 or SRF KD were most enriched in the same subsets (Figure 5C), indicating that the TGFβ-NNMT-ATF4/SOX9/SRF network operates within the disease-expanded myofibroblast lineage.

**Figure 5.**
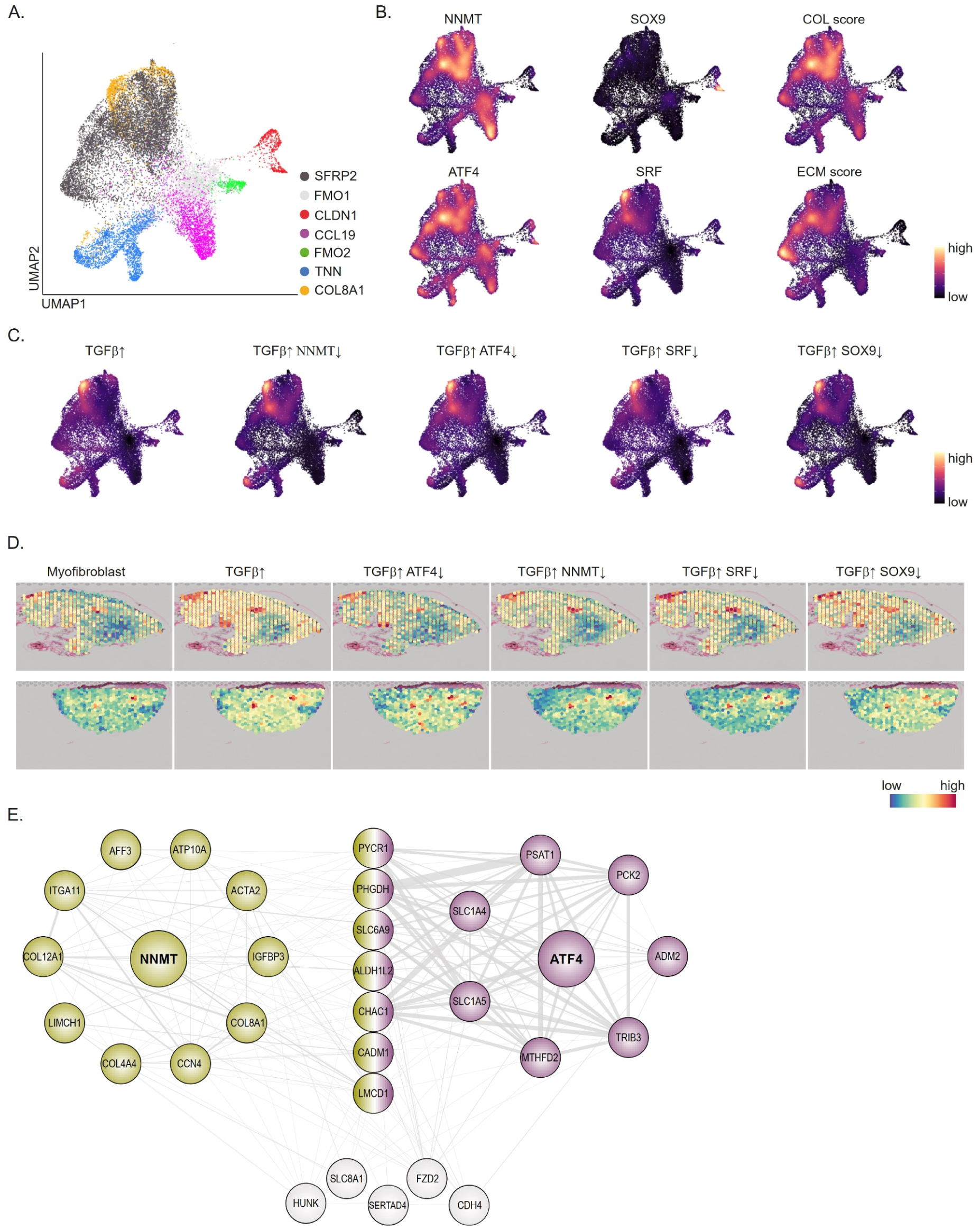
The TGFβ-NNMT-ATF4/SOX9/SRF network localizes to disease-expanded myofibroblasts lineage and niches in SSc. **A.** UMAP of dermal fibroblasts (DF) subtypes in scRNA-Seq data from SSc skin (23). **B.** UMAP feature plots showing expression of NNMT, ATF4, SOX9 and SRF, together with collagen (Col) and extracellular (ECM) module scores. **C.** Integration of RNA-Seq data from TGFβ-stimulated ATF4, NNMT, SOX9 and SRF knockdown (KD) experiments. TGFβ-upregulated and KD-attenuated gene sets were projected onto the scRNA-Seq dataset shown in panel A. **D.** Spatial transcriptomic maps from publicly available SSc datasets (23) showing localization of a myofibroblast gene score, TGFβ-upregulated genes, and KD attenuated-genes sets. **E.** Co-expression network analysis reveals potential gene targets of NNMT and ATF4. The thickness of the edges represents the cumulative weight of all interaction for the corresponding node pair.

We next examined the spatial context of this network using publicly available spatial transcriptomic data from SSc skin. Myofibroblasts showed broad co-localization with TGFβ-upregulated genes as well as with NNMT-, ATF4-, SOX9-, and SRF-dependent TGFβ subsets (Figure 5D), reinforcing that this regulatory axis underpins ECM-rich, myofibroblast-dominated dermal niches.

To identify putative genes directly co-regulated by NNMT, ATF4, SOX9, and SRF, we performed a co-expression analysis and constructed a network of candidate genes based on known interactions. The NNMT-containing co-expression module included ATF4, but not SOX9 or SRF, suggesting a stronger co-regulatory relationship between these two. Notably, both NNMT and ATF4 converged within a highly specific shared network, with NNMT showing potential interactions with ECM components (COL8A1, COL4A4, COL12A1, ITGA11) and fibroblast activation markers (ACTA2), whereas ATF4 was linked to genes involved in metabolic adaption and amino acid transports (SLC1A5, SLC6A9, PCK2, ALDH1L2).

Together, these analyses position the NNMT-dependent ATF4/SOX9/SRF network within the SFRP2+/COL8A1+ myofibroblast lineage in human skin, linking the in vitro functional dependencies of these factors to the in vivo cellular and spatial contexts where profibrotic SSc programs are executed.

## Discussion

Progressive skin fibrosis is the hallmark and delineates disease subsets. The sclerosing process may result profound disfigurement, functional limitations and complications such as contractures with loss of hand function, restricted mobility, microstomia, and digital ulcers that often resist treatment and may culminate in amputation. The extent of skin fibrosis further correlates with internal organ involvement, particularly interstitial lung disease, which remains the leading cause of SSc-related mortality (1). Despite recent therapeutic advances targeting inflammation, current treatment strategies fail to directly address the aberrant activation of DFs, the central effectors of cutaneous fibrosis. This lack of DF-directed strategies underscores the urgent need for novel molecular targets within this compartment (3, 6).

Our study identifies NNMT as a key metabolic regulator of DF activation in SSc. NNMT was highly expressed in SSc skin across multiple transcriptomic datasets, particularly within disease-expanded SFRP2+/COL8A1+ fibroblast populations and ECM-rich dermal niches, linking its expression directly to profibrotic fibroblast states. Functional studies revealed that NNMT depletion increased the SAM/SAH ratio, restored levels of H3K27me3, a histone mark previously linked to DF activation (36), and suppressed ECM programs, collagen organization, and cytoskeletal remodeling.

Mechanistically, we demonstrate that NNMT is TGFβ-inducible and required for the full activation of TGFβ-dependent transcriptional programs. NNMT loss attenuated TGFβ-induced ECM gene expression and COL1A1 secretion. Moreover, NNMT was necessary for TGFβ-driven induction of the TFs ATF4, SOX9, and SRF, which together form a convergent network controlling ECM organization and myofibroblast differentiation. Notably, ATF4 and SOX9 have been implicated in fibrotic processes across multiple organs, including lung (37–39), liver (40), kidney (41), and heart (42, 43), highlighting their relevance as profibrotic transcriptional regulators. SRF, while less directly studied in fibrosis, regulates cytoskeletal and contractile gene expression, consistent with a role in myofibroblast activation. The TGFβ–NNMT–ATF4/SOX9/SRF axis therefore represents a previously unrecognized regulatory circuit that integrates metabolic and transcriptional pathways in SSc fibroblasts. Single-cell and spatial transcriptomic analyses localized this network to myofibroblast-rich dermal regions, emphasizing its in vivo relevance to fibrotic remodeling.

Interestingly, NNMT was also highly expressed in CCL19+ DFs, a disease-adapted subset previously shown to engage in crosstalk with immune cells (33). These DFs co-expressed ATF4 but lacked SOX9, SRF, and TGFβ signature genes, suggesting that NNMT may support DF activation programs independent of TGFβ signalling. The transcriptional profile of CCL19+ DFs, characterized by immune-interacting features, implies that NNMT’s role may extend beyond disease-associated myofibroblasts, linking it to immune-associated DF populations that shape the inflammatory-fibrotic interface in SSc.

While we did not directly dissect how NNMT regulates the expression of these TFs, it likely acts through modulation of cellular methylation potential. Several studies have shown that NNMT abundance is tightly linked to the SAM/SAH balance, which shapes epigenetic regulation of gene expression (9). Thus, NNMT represents a unique metabolic node coupling cellular metabolism to transcriptional control. A similar mechanism has been demonstrated in CAFs (15, 35), supporting the notion that NNMT-dependent metabolic reprogramming constitutes a conserved strategy underpinning fibroblast activation across diverse pathological settings.

The broader relevance of NNMT to fibroblast activation is reinforced by evidence from other diseases. NNMT is a hallmark of CAFs, where it promotes matrix remodeling and tumor desmoplasia (15). Similarly, NNMT upregulation has been reported in idiopathic pulmonary fibrosis (44), as well as in hepatic (10) and renal fibrosis (16), where its inhibition mitigated tissue remodeling. Collectively these findings point to a conserved profibrotic function of NNMT across organs, although some studies suggest context-dependent roles, particularly in the kidney (45). The convergence of evidence from cancer, metabolic disease, and organ fibrosis indicates that NNMT participates in a shared fibroblast activation program rather than a tissue-restricted process. Consistent with this concept, our data show that NNMT is induced by TGFβ and required for TGFβ-driven profibrotic gene expression in DFs, suggesting a unifying mechanism by which profibrotic stimuli co-opt NNMT activity to sustain fibroblast activation.

Our data suggest that NNMT may represent a promising therapeutic target for SSc. Small-molecule NNMT inhibitors are under active development, notably a potent and selective compound recently shown to reduce tumor burden and metastasis in multiple mouse models of cancer by modulating CAFs (35). If such inhibitors can be repurposed or optimized for fibrotic indications, they could represent the first class of therapies directly targeting DF activation in SSc. Although our current study is limited to in vitro models and siRNA-mediated KD, the consistency of findings across publicly available datasets, and DF cultures provides a compelling rationale for advancing to preclinical testing in SSc relevant models. Future research should validate NNMT inhibition in vivo assessing its effects on both cutaneous and internal organ fibrosis.

In summary, we identify NNMT as a previously underappreciated driver of DF activation and dermal fibrosis in SSc. By linking TGFβ signaling to metabolic control of transcription, NNMT sustains profibrotic gene expression and ECM production through an ATF4/SOX9/SRF-dependent network. Together with prior evidence from cancer biology and organ fibrosis, these data position NNMT as a shared fibroblast effector and a druggable candidate for antifibrotic therapy. Targeting NNMT may therefore represent a novel and urgently needed strategy to halt or reverse tissue fibrosis in SSc.

## Acknowledgments

We thank the BSF for supporting the RNA-Seq and the patients and clinical staff of the Medical University of Vienna for their participation. We are also very grateful to H. S. Afarani for her valuable assistance with experimental work. The Vienna BioCenter Core Facilities (VBCF) Metabolomics Facility acknowledges funding from the Austrian Federal Ministry of Education, Science & Research; and the City of Vienna.

## Author contributions

Conceptualization: AT, TK

Methodology: AT, KvD, AK, TKö, LXH, TK.

Investigation: KvD, FK, BN, LXH, TKö.

Formal analysis: AT, FK, TK.

Visualization: AT, FK, JMS, TK.

Resources: PH, BL, DA.

Writing – original draft: TK, AT.

Writing – review & editing: all authors.

Supervision: TK.

## Conflict of Interest Statement

D.A. reports no conflicts of interest related to this work. For transparency, D.A. discloses receiving grants from Eli Lilly and speaker or consultancy fees from AbbVie, Gilead, Johnson & Johnson, Eli Lilly, MSD, Novartis, and Sandoz. The other authors declare no conflicts of interest.

