## Supplementary Figures for "Nicotinamide N-Methyltransferase drives fibroblast activation and skin fibrosis in systemic sclerosis"

[3] Vienna BioCenter Core Facilities, Vienna, Austria.

*Corresponding author

**Correspondence to:**

Assoc. Prof. Dr. Thomas Karonitsch

Division of Rheumatology, Department of Medicine 3

Medical University of Vienna

Spitalgasse 23

1090 Vienna, Austria

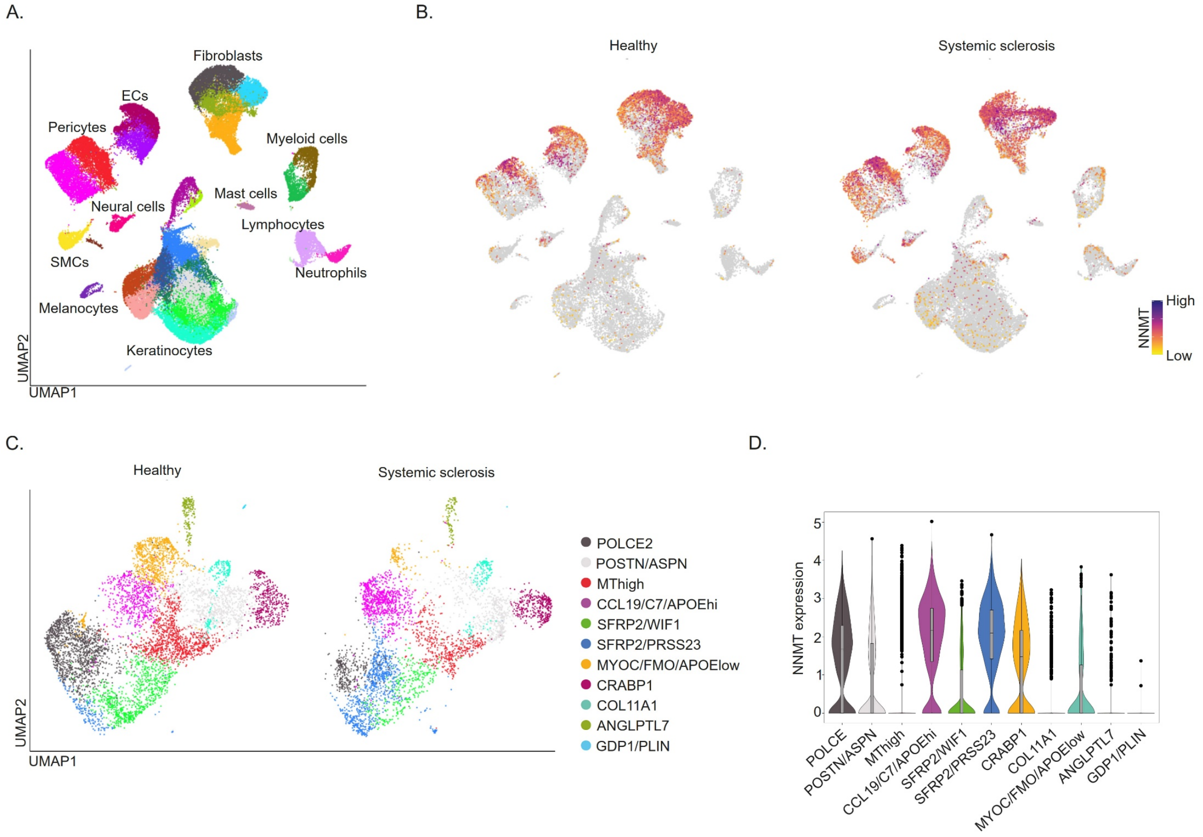


**Supplementary Figure 1.**

**A.-D.** Analysis of publicly available scRNA-Seq data from skin biopsies of SSc (n=12) patients and HCs (n=10) (22).

**B.** UMAP plot showing major skin cell populations.

**C.** UMAP plots showing NNMT expression across cell types in HC and SSc.

**D.** UMAP plot showing DF subtypes identified in HC and SSc.

**E.** Violin plots showing NNMT expression levels across DF subtypes.

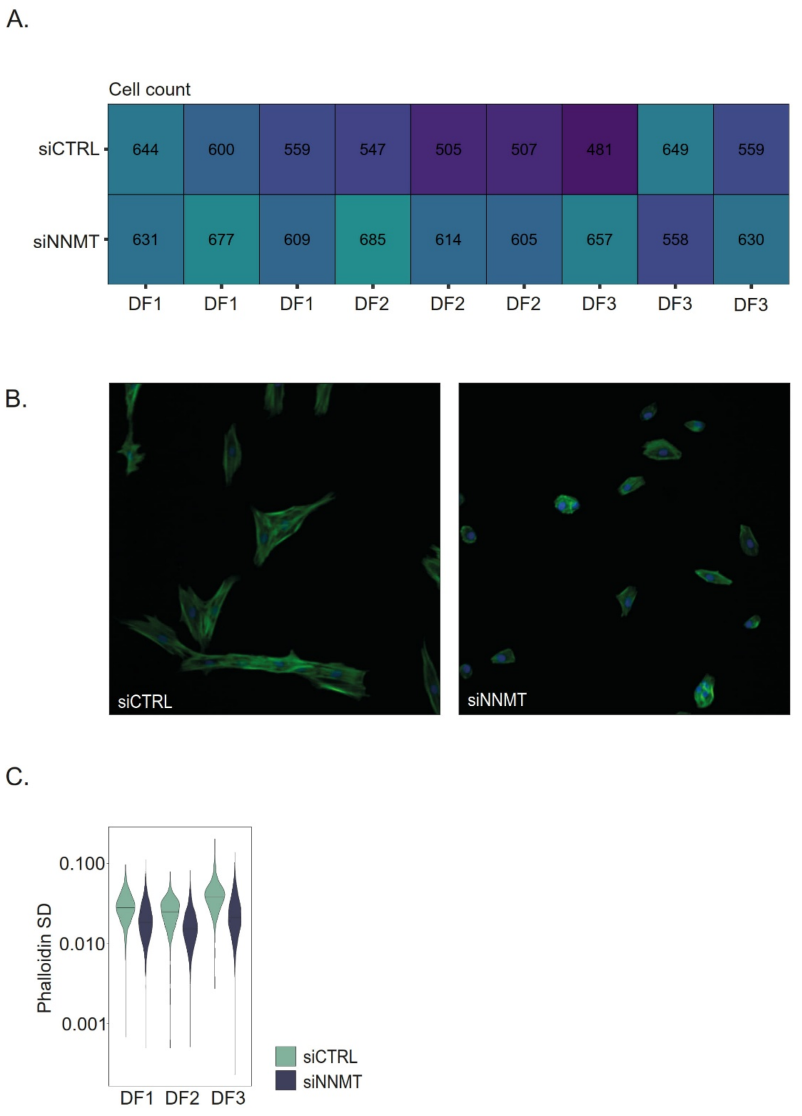


**Supplementary Figure 2.**

**A-C.** DFs from three different donors (DF1, DF2 and DF3) were stained with DAPI and phalloidin after transfection with non-targeting control or NNMT-targeting siRNA pools. Images were acquired using a PerkinElmer Opera microscope and analyzed with CellProfiler software.

**A.** Cell counts of DFs seeded in 96-well plates and cultured for 24 hours.

**B.** Representative images of DFs stained with DAPI and phalloidin.

**C.** Violin plots showing the standard deviation (SD) of phalloidin intensity as a measure of cytoskeletal organization.

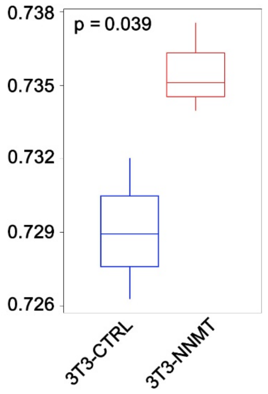


**Supplementary Figure 3.**
Reanalysis of publicly available RNA-Seq data from 3T3 cells overexpressing NNMT and corresponding control cells (3T3-CTRL) (15). NNMT-downregulated genes in DFs identified in our NNMT knockdown (KD) RNA-Seq experiments (Figure 2) were assessed for their expression in the 3T3 dataset. Box plots show that the expression of NNMT-downregulated genes in DFs was significantly upregulated in 3T3-NNMT cells.

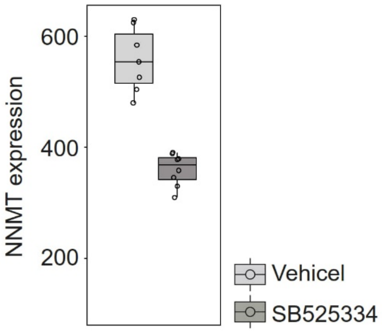


**Supplementary Figure 4.**
NNMT mRNA expression in publicly available bulk RNA-Seq data (20) from skin samples from bleomycin-treated mice receiving TGFβ receptor inhibitor SB525334 or vehicle control.

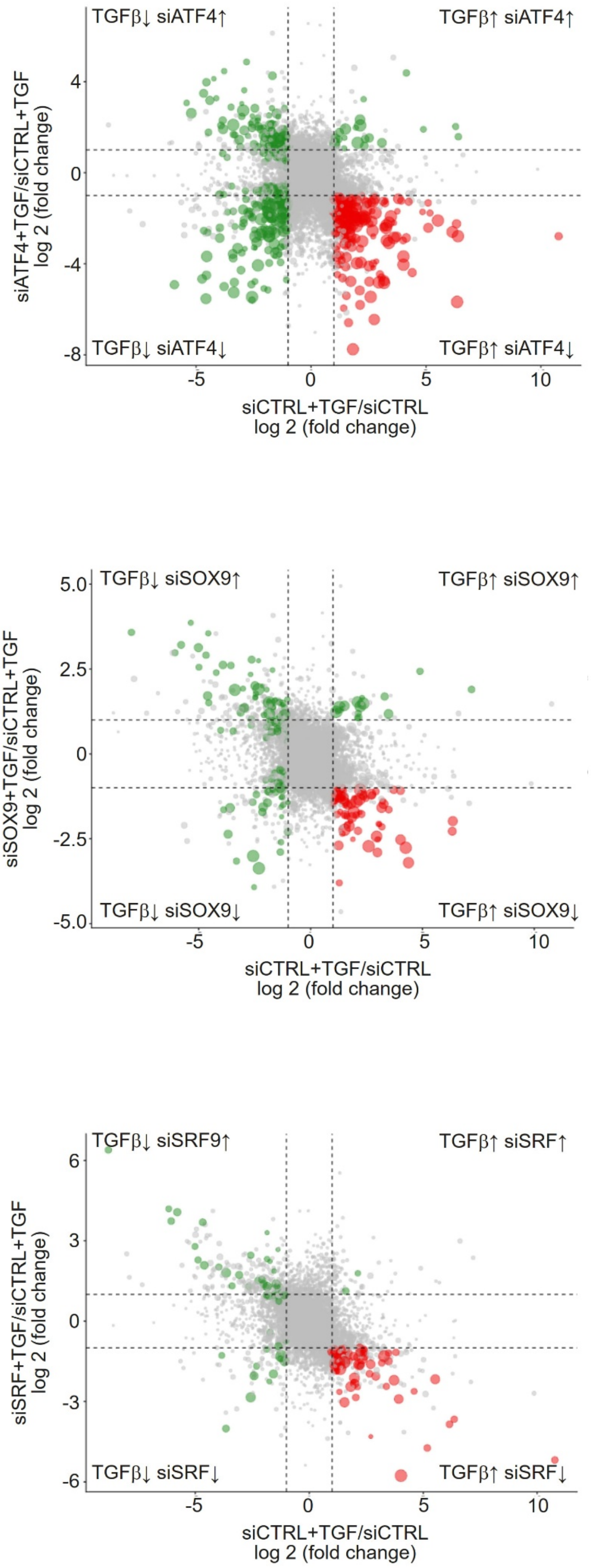


**Supplementary Figure 5.**
Scatter plot illustrating the influence of ATF4, SOX9 and SRF knockdown (KD) on TGFβ-regulated genes, as measured by RNA-Seq (n=5). Genes highlighted in red (lower right quadrant) represent TGFβ-induced genes whose induction is blunted by the KD. n refers to the number of independent donors (biological replicates).
